# Identification and functional validation of an enhancer variant in the 9p21.3 locus associated with glaucoma risk and elevated expression of *p16^INK4a^*

**DOI:** 10.1101/2023.05.18.541339

**Authors:** Yizhou Zhu, Cagdas Tazearslan, Michael G. Rosenfeld, Andras Fiser, Yousin Suh

**Affiliations:** Department of Obstetrics and Gynecology, Columbia University, New York, NY10032, USA; Department of Genetics, Albert Einstein College of Medicine, Bronx, NY10461, USA; Department of Medicine, School of Medicine, University of California, San Diego, La Jolla, CA 92093, USA; Howard Hughes Medical Institute, University of California, San Diego, La Jolla, CA 92093; Department of Systems & Computational Biology, Albert Einstein College of Medicine, Bronx, New York 10461, USA; Department of Biochemistry, Albert Einstein College of Medicine, Bronx, New York 10461, USA; Department of Genetics and Development, Columbia University, New York, NY10032, USA

**Keywords:** Glaucoma, YY1, p16INK4A, 9p21, Genome-wide Association Study, Cellular Senescence, Molecular Genetics

## Abstract

Glaucoma is a leading cause of irreversible blindness, with advanced age being the single most significant risk factor. However, the mechanisms underlying the relationship between aging and glaucoma remain unclear. Genome-wide association studies (GWAS) have successfully identified genetic variants strongly associated with increased glaucoma risk. Understanding how these variants function in pathogenesis is crucial for translating genetic associations into molecular mechanisms and, ultimately, clinical applications. The chromosome 9p21.3 locus is among the most replicated glaucoma risk loci discovered by GWAS. Nonetheless, the absence of protein-coding genes in the locus makes interpreting the disease association challenging, leaving the causal variant and molecular mechanism elusive. In this study, we report the identification of a functional glaucoma risk variant, rs6475604. By employing computational and experimental methods, we demonstrated that rs6475604 resides in a repressive regulatory element. Risk allele of rs6475604 disrupts the binding of YY1, a transcription factor known to repress the expression of a neighboring gene in 9p21.3, p16INK4A, which plays a crucial role in cellular senescence and aging. These findings suggest that the glaucoma disease variant contributes to accelerated senescence, providing a molecular link between glaucoma risk and an essential cellular mechanism for human aging.

## Main Text

Glaucoma is a group of optic neurodegenerative diseases that lead to progressive loss of retinal ganglion cells (RGCs)^1,2^. In the United States and worldwide, glaucoma has emerged as a leading cause of irreversible blindness among older individuals^3^. Numerous studies have identified older age as the most significant and consistent risk factor for glaucoma development^4-8^. Despite this correlation, the precise mechanisms through which age-related pathophysiological changes in RGCs increase disease susceptibility remain largely unknown.

Genome-wide association studies (GWAS) have consistently identified the 9p21.3 locus as a critical risk factor for glaucoma across various populations^9-12^. The precise role of this approximately 150 kb non-coding region in disease risk, however, remains unclear. Epigenetic annotations have uncovered numerous putative regulatory elements within the 9p21.3 locus, pointing to its possible regulatory functions in relation to nearby genes^13^. The INK4 family genes, including CDKN2A (encoding p16^INK4A^ and p14^ARF^) and CDKN2B (encoding p15^INK4B^), which are situated immediately upstream of the locus, play a crucial role in cellular senescence and are considered reliable aging markers in both mice and humans^14,15^.

Recent studies have demonstrated that increased p16^INK4A^ expression can lead to RGC senescence in cell culture, animal models, and human glaucoma retinas^16^. Based on these findings, we propose that the non-coding glaucoma risk variants within the 9p21.3 locus may have regulatory functions for p16^INK4A^ and could contribute to glaucoma risk by modulating its expression. Unraveling the complex relationship between the 9p21.3 locus, the INK4 family genes, and RGC senescence could provide valuable insights into the underlying mechanisms of age-related glaucoma and pave the way for the development of novel therapeutic approaches to combat this debilitating disease.

To identify functional variants associated with glaucoma risk in 9p21.3, we obtained the complete list of available glaucoma GWAS from the NHGRI catalogue (**Table S1**). Six independent studies, including three European and three Japanese populations, have reported six glaucoma-associated common variants in the 9p21.3 locus. Due to strong linkage disequilibrium (LD) in 9p21.3, multiple variants in the locus were found as significantly associated with glaucoma. Furthermore, the functional variant can be one of the reported GWAS index variants or one in high LD with them. To list the possible candidate functional variants, we examined the LD patterns of the locus using genotyping data from the 1000 Genomes Project. This resulted in an additional 40 candidate variants that were tightly linked (r^2^ > 0.8) to the glaucoma risk variants (**Figure 1A**). Besides rs1063192, a 3’ UTR variant for CDKN2B, all other variants resided in the intronic region of CDKN2B-AS1, a long non-coding RNA in the 9p21.3 locus.

**Figure 1.**
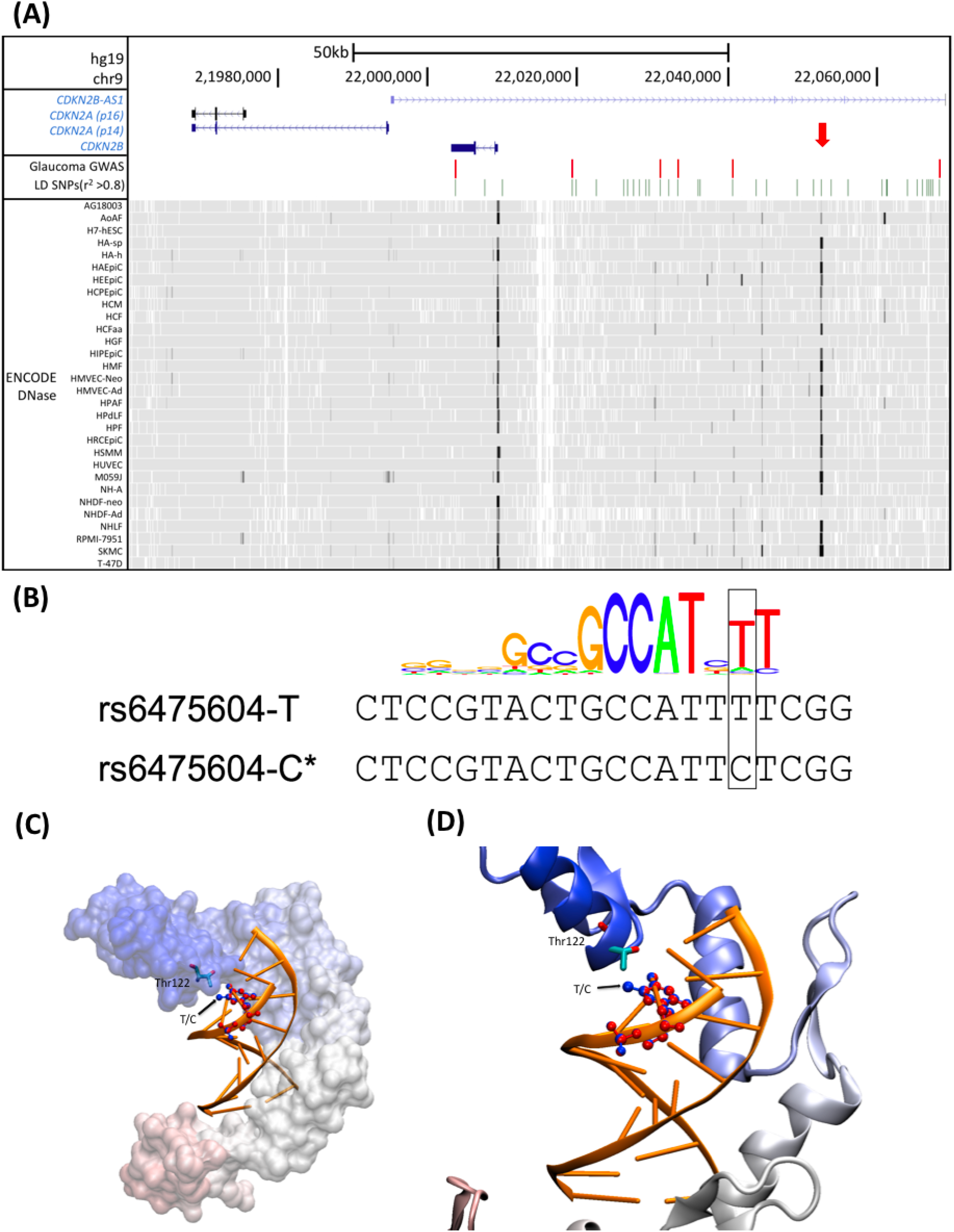
(**A**) Overview of the 9p21.3 glaucoma risk locus. Locations of glaucoma GWAS risk variants and linked (r^2^ > 0.8) variants (LD SNPs) are indicated in red and green bars, respectively. The bottom DNase hypersensitivity panel from the ENCODE Project suggests putative enhancers in this locus. Red arrow indicates the conserved enhancer that harbors the functional risk variant, rs6475604. (**B**) Overlay between the consensus motif of human YY1 and the predicted YY1 motif at rs6475604. The glaucoma risk allele (C) disrupts a conserved T nucleotide. **(C)** Superimposed complex structures of YY1 with two alternative DNA fragments bound to it. The protein model is shown in surface model, colored from red to blue from the N to C terminus. For clarity the surface model is transparent. The superposed DNA fragments are shown in ribbon models (orange). The only different position T/C is shown in CPK models, blue corresponds to T and red to C nucleic acid base (black arrow). The coordinating Thr122 residue in the YY1 protein is shown in stick model with element based coloring. **(D)** A close up of panel C, showing the coordinating Thr122 from YY1 in stick model and the nucleic acid base residues in blue (T) and in red (C). Figures were prepared with VMD program.

To narrow down the list of candidate causal variants, we examined epigenomic data from the ENCODE and Roadmap Projects to screen for variants that resided in putative regulatory elements. We found that the 9p21.3 locus was enriched in enhancers, the principal regulatory components of the genome that regulate transcription over a long distance^17^. By interrogating the DNase hypersensitive sites for the entire glaucoma LD block (the minimum chromosomal segment containing all LD variants), we found several cell type-specific weak enhancers, as well as one strong and well-conserved enhancer that was present in most of the cell lines (**Figure 1A**). To further consolidate our *in silico* regulatory function prediction, we annotated the candidate glaucoma risk variants using RegulomeDB, a database that annotates common variants with known and predicted regulatory elements^18^. RegulomeDB ranks variants from 1 (most likely) through 6 (least likely) to affect regulatory function. Among all, rs6475604 was the highest-ranked variant with a score of 2a (**Table S2**), which we found to be uniquely located in the identified conserved enhancer (**Figure 1A**).

The variant rs6475604 is a C/T polymorphism with the C allele associated with the risk of glaucoma. RegulomeDB predicts that rs6475604 is in the canonical and actual binding site of a transcription factor, YY1, and the risk rs6475604-C allele would interrupt the consensus binding site of YY1 (**Figure 1B**). We further performed computational modeling of the YY1 protein with both the risk and non-risk allele. We explored all possible 9-mer DNA fragments in complex with the YY1 transcription factor using the TF2DNA modeling tool, which uses a structure-based affinity calculation^19^. When ordered by relative binding affinities, the two motifs containing the risk and non-risk allele (CGCCATT<u>C</u>T and CGCCATT<u>T</u>T) appeared as the 66th and 250th strongest binding DNA fragment among the 262,144 possible combinations (4^9^), indicating that these two fragments were among the most likely binding site candidates (among the top 0.1% of all). Furthermore, the segments CGCCATT<u>T</u>T and CGCCATT<u>C</u>T had relative binding affinities of 487.21 and 465.71, respectively (the higher values indicate stronger binders), demonstrating that the glaucoma risk fragment had reduced binding of YY1 as compared to the non-risk allele. The two segments differ only by the C/T change in position 8, which is a single methyl group difference in their pyrimidine rings. In the structural model, this extra methyl group in T, the non-risk allele, appears to interact with the methyl group of Thr122 in the YY1 protein (**Figure 1C and D**), providing an extra van der Waals interaction and a better overall binding pattern.

Interestingly, YY1 is a general transcription factor known to function as a repressor of p16^INK4A^ gene expression^20^ and to interact with critical negative regulators of p16^INK4A^, such as polycomb complexes^21^. If the computational prediction is accurate, the glaucoma risk allele (rs6475604-C) would increase p16^INK4A^ expression by causing reduced binding of its repressor, YY1. To experimentally test the predicted disruption of YY1 binding by rs6475604-C compared to the non-risk rs6475604-T allele, we performed a competitive gel mobility shift assay (EMSA) using the long YY1 consensus motif (GCCGCCATTTTG) as a biotin-labeled probe^22^ and the predicted YY1 motif at rs6475604 as unlabeled oligonucleotide competitors for binding of recombinant YY1 (**Fig S1**). We found that the non-risk allele (T)-containing oligonucleotide was a much more effective competitor compared to the risk allele (C), leading to a significant reduction in the intensity of the YY1-probe complex, especially when applied at 5x concentration (**Figure 2A**). Notably, the YY1 consensus motif itself (unlabeled probe) showed complete competition under this condition. Reciprocally, we performed the same assay using the predicted YY1 motif containing either the non-risk or risk allele as labeled probes. The result was consistent, showing that the non-risk allele competed better for binding of YY1 regardless of the probes used (**Figure 2A**).

**Figure 2.**
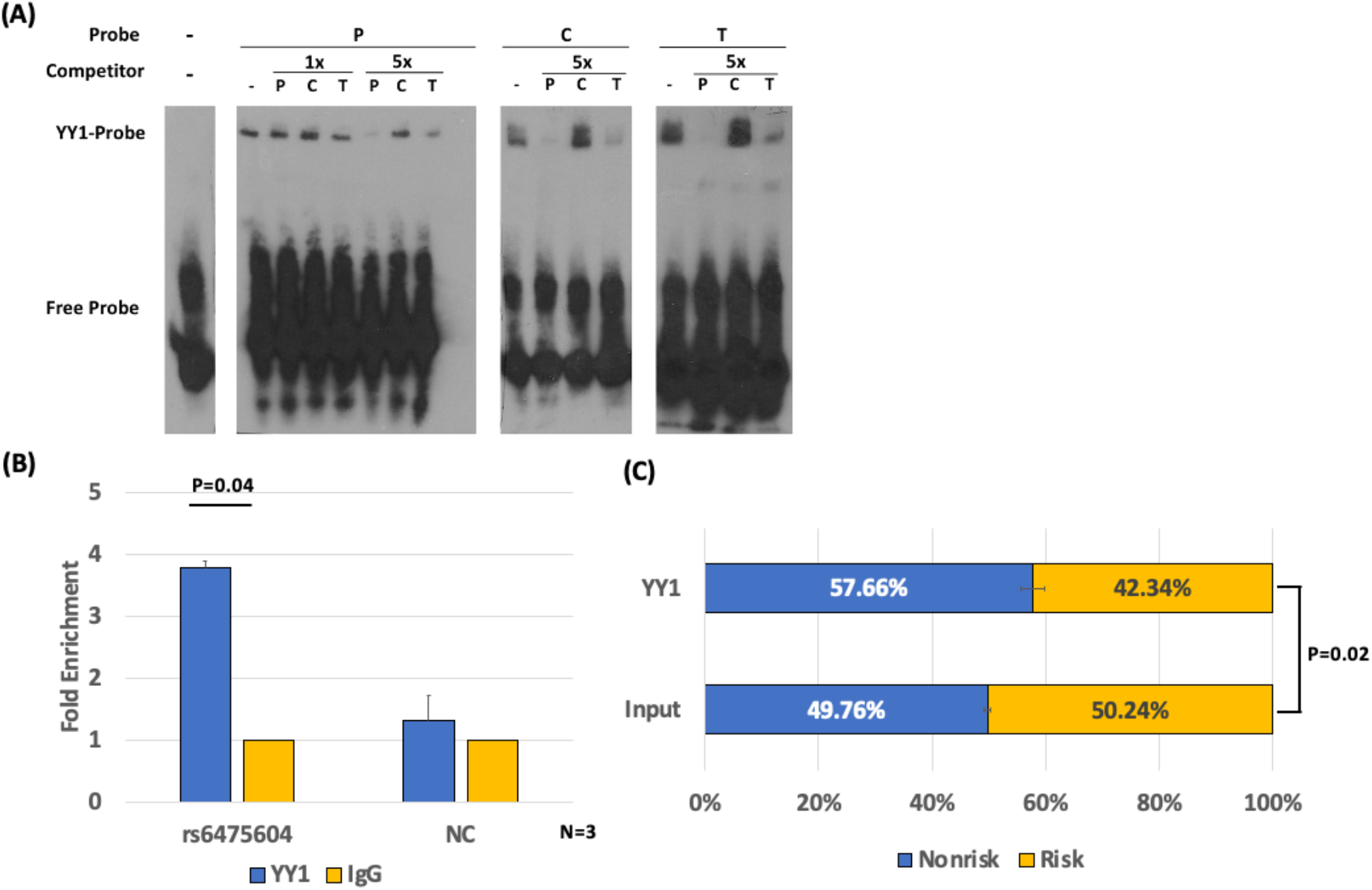
**(A)** Electrophoretic mobility shift affinity (EMSA) assays of YY1 with rs6475604. P: YY1 consensus motif; C: rs6475604-C allele; T: rs6475604-T allele. Binding was performed under no competition (-) or with unlabeled competitors in 1x or 5x concentrations. **(B)** ChIP-qPCR and **(C)** allele-specific digital PCR of YY1 binding on rs6475604 in neural stem cells differentiated from human embryonic stem cell (H13) (N=3). Negative control (NC) was a nearby locus with no marks of enhancer YY1 binding from ENCODE data. Error bars indicate SEM. Statistics was performed using two-sided t test.

To validate the EMSA result in vivo, we performed ChIP-qPCR in neural stem cells differentiated from H13 embryonic stem cells. We found that endogenous YY1 significantly bound to the variant locus (3.8-fold enrichment, **Figure 2B**). To determine the impact of rs6475604 on YY1 binding, we utilized the heterozygote genotype of the variant in H13 and performed allele-specific digital PCR, which indicated a significant allelic imbalance of the non-risk allele over the risk allele (**Figure 2C**). These observations demonstrated that the glaucoma risk allele (rs6475604-C) significantly diminished YY1 binding compared to the non-risk allele (rs6475604-T), consistent with the in vitro results.

To investigate the functional outcome of the YY1 binding alteration caused by rs6475604, we performed a luciferase reporter assay for the enhancer harboring the variant in four cell lines (**Figure S2A**). We found that the putative enhancer significantly down-regulated the reporter in HeLa cells, while showing insignificant trends of down-regulation in IMR90, MCF7, and HEK293. A comparison of the reporter harboring different alleles of rs6475604 showed that the glaucoma non-risk allele was consistently associated with stronger reporter repression in all cell lines, although the difference was subtle and insignificant. Taken together, the results suggested that while the rs6475604-harboring enhancer is likely a repressive regulatory element, its function is cell type and context-dependent.

Lastly, utilizing the linked glaucoma variant (rs1063192) in the 3’UTR of CDKN2B, we performed allele-specific qPCR to test whether the alteration of YY1 binding by rs6475604 would affect p15^INK4B^ expression. In H13-derived NSCs and HEK293, both of which harbored heterozygote 9p21.3 glaucoma genotypes, we did not identify an imbalance between risk and nonrisk alleles (**Figure S2B**), suggesting that the gene was likely not a target.

In this study, we uncovered a functional variant, rs6475604, located in an enhancer region of the 9p21.3 locus by analyzing and integrating data from GWAS, epigenomics, computational modeling, and experimental validation. Our findings revealed that this enhancer variant led to decreased binding of the YY1 transcription factor, a known repressor of p16INK4A gene expression. This observation aligns with previous studies that demonstrated the recruitment of YY1 to the INK4A-ARF gene body, along with other polycomb group proteins such as BMI-1 and EZH2, which negatively regulate p16^INK4A^ expression^20,23,24^.

The 9p21.3 non-coding region has been linked to various age-related disorders apart from glaucoma, including cancers, coronary artery diseases (CAD), and type 2 diabetes (T2D). Fine-mapping of risk variants has suggested that the causal mechanisms underlying each disease are likely distinct ^15^. For CAD, several studies have pointed to p15^INK4B^ or its antisense long non-coding RNA CDKN2B-AS1 as the causal effector^13,25,26^. In contrast, our research indicates that the expression of p15^INK4B^ remains unaltered by the genotype of the glaucoma risk variant, thus presenting p16^INK4A^ as an alternative effector gene.

High expression levels of p16^INK4A^ are crucial for inducing cellular senescence^27,28^. As organisms age, senescent cells accumulate and contribute to numerous age-related diseases such as osteoarthritis, cataracts, tumorigenesis, cardiac hypertrophy, renal dysfunction, lipodystrophy, and sarcopenia in mice^29^. Previous studies have reported increased p16^INK4A^ expression in glaucomatous eyes in mice and humans, with elevated p16^INK4A^ levels being causally connected to RGC senescence in mouse models^16^. Furthermore, the glaucoma risk genotype in SIX6, an activator of p16^INK4A^ expression, has been associated with higher p16^INK4A^ expression and senescence in human RGCs^16^. In light of these findings, our data suggest that genetic variants that increase p16^INK4A^ expression may contribute to glaucoma risk by enhancing the propensity for cellular senescence, a fundamental aging mechanism, in humans. This understanding could potentially inform future therapeutic strategies targeting cellular senescence in glaucoma diseases.

### Experimental Procedures

#### SNP selection

SNP Selection The list of glaucoma-associated GWAS variants was obtained from the GWAS Catalogue (https://www.ebi.ac.uk/gwas/, as of November 2017), a comprehensive database for GWAS that currently includes 2,854 publications and 33,674 unique SNP-trait associations. The default filter for p-value (p<5×10^-8^) was used to search for significant associations, and no additional filter on effect size was applied. Variants were considered in the 9p21.3 locus if they were located within the coordinates, chr9:21950000-22180000 (hg38).

### Linkage Disequilibrium (LD) analysis

We used SNAP (SNP Annotation and Proxy Search) to examine the reported GWAS variants to obtain a full list of SNPs in LD with the 9p21 glaucoma variants (Johnson et al. 2008). Variants with LD r^2^ >= 0.8 were included for downstream analysis of in silico function prediction. The variants found in the 1000 Genomes and HapMap Projects were combined to obtain the final list, after removing redundancies. For determining the risk allele for LD variants, we constructed haplotypes using Haploview (Barrett et al. 2005) and the individual genotype information of the 1000 Genomes Project.

### RegulomeDB

RegulomeDB is a bioinformatics tool for predicting the regulatory function of ∼60 million annotated variants (Boyle et al. 2012). The database is built upon 962 epigenomic datasets from the ENCODE consortium, NCBI Sequence Read Archive, and other sources, including a large collection of eQTL studies. The database ranks variants from 1 (highest) to 6 (lowest) for their likelihood of exerting regulatory functions. Rank 1 and 2 variants are considered likely to be functional based on strong evidence from multiple datasets. The tool is freely accessible at http://www.regulomedb.org. We applied RegulomeDB to all 9p21 glaucoma variants and GWAS variants to examine potential regulatory functions.

### Electrophoresis mobility shift affinity (EMSA) assay

The following sequences were used for the gel mobility shift assay:

YY1-consensus: CGCTCCGT**GCCGCCATTTTG**GGCGGCTGGT

rs6475604-C[Risk]: CGCTCCGT**ACTGCCATTCTC**GGCGGCTGGT

rs6475604-T[Non-Risk]: CGCTCCGT**ACTGCCATTTTC**GGCGGCTGGT

The 3’-biotinylated probes and unlabeled competitors were synthesized by IDT and annealed with the corresponding reverse complements. Forty nanograms of probes were mixed with 200 ng recombinant YY1 (31332, Active Motif) in a buffer containing 10 mM Tris-HCl (pH 7.4), 150 mM KCl, 0.1 mM DTT, and 0.1 mM EDTA, and incubated for 20 min at room temperature. For competition binding assays, unlabeled competitors with 1X and 5X concentrations were pre-incubated with YY1 protein before adding labeled probes. The incubated protein-DNA complex was loaded on a 6% polyacrylamide gel and run at 100 V for 50 min in 1x TBE buffer. The gel was then transferred to a nitrocellulose membrane and probed with streptavidin-HRP (Abcam ab7403) and imaged on exposure film (Thermo 34090) using ECL substrate(Thermo 32106).

### *In silico* calculation of relative binding affinities

Relative binding affinities were obtained using the TF2DNA algorithm (Pujato et al. 2014). Briefly, TF2DNA construct a three dimensional computational homology model of the complex of the protein transcription factor and cognate DNA fragment and assesses the relative binding affinity using a knowledge-based statistical pair potential, after optimization of the complex in a molecular mechanics force field. In case of YY1 transcription factor 9 residue long, double stranded DNA fragments were explored in all possible combinatorial variation, resulting in (49)=262,144 different complexes and corresponding energies.

### Cell Culture

Human embryonic stem cells H13 (WiCell WA13) were maintained in a feeder-free E8 system (ThermoFisher A1517001). To differentiate into neural stem cells, ES cells were sub-cultured at 15% confluency, and cultured in PSC Neural Induction Medium (Thermo A1647801) for 6 days as the manufacturer indicated. After reaching confluency, cells were maintained in neural expansion medium (49% Neurobasal medium, 49% Advanced DMEM-F12, 2% neural induction supplement) for two passages before collection.

### Allele-Specific digital PCR

RNA was isolated using the PureLink RNA Mini Kit (ThermoFisher) according to the manufacturer’s instructions. DNase treatment was performed using the DNA-free DNase Treatment & Removal Kit (ThermoFisher). To synthesize cDNA, reverse transcription was performed using SuperScript IV Reverse Transcriptase (ThermoFisher) with oligo-dT primers, following the manufacturer’s instructions.

To perform allele-specific expression analyses of CDKN2B, 50 ng of cDNA was loaded onto the Quantstudio 3D digital PCR system (version 2, ThermoFisher) and amplified using Taqman SNP arrays (rs1063192, ThermoFisher). Allele-specific signal quantification was performed using the online cloud application provided by the manufacturer.

### Chromatin immunoprecipitation

For crosslinking, 10 million NSCs were crosslinked using 1% formaldehyde (Thermo 28908) in PBS for 10 min. The reaction was quenched by incubating in 125 mM glycine for 5 min, and cells were washed with PBS twice and harvested using a silicone scraper. To create nuclear lysate, cells were resuspended sequentially in 10 mL Buffer 1 (50 mM HEPES-KOH pH 7.5, 10 mM NaCl, 1 mM EDTA, 10% glycerol, 0.5% IGEPAL CA-630, 0.25% Triton X-100), 10 mL Buffer 2 (10 mM Tris-HCl, pH 8.0, 200 mM NaCl, 1 mM EDTA, 0.5 mM EGTA), and 900 μl Buffer 3 (10 mM Tris-HCl, pH 8.0, 100 mM NaCl, 1 mM EDTA, 0.5 mM EGTA, 0.1% Na-Deoxycholate, 0.5% N-lauroylsarcosine). In Buffer 3, the lysate was sonicated using a Bioruptor (Diagenode, 12 min at high energy output, 30 sec on, 30 sec off), and mixed with 10 μg YY1 antibody (Santa Cruz sc-7341) or IgG (Thermo 10400C) in 100 μl Protein G beads (Thermo 10003D) for 14 hr immunoprecipitation at 4 ^o^C. The beads were cleaned up with 2x washes of low salt buffer (20 mM Tris-HCl, pH 7.5, 150 mM NaCl, 2 mM EDTA, 0.1% SDS, 1% Triton-X100), 2x of high salt buffer (20 mM Tris-HCl, pH 7.5, 500 mM NaCl, 2 mM EDTA, 0.1% SDS, 1% Triton-X100), and 1x LiCl buffer (10 mM Tris-HCl, pH 7.5, 250 mM LiCl, 1 mM EDTA, 1% Na-Deoxycholate, 1% IGEPAL CA-630). The bead-bound complex was eluted with 200 μl elution buffer (50 mM Tris-HCl, pH 8.0, 10 mM EDTA, 1% SDS) for 30 min at 65°C. The eluent was sequentially treated with 0.2 mg/mL RNase A and 0.2 mg/mL Proteinase K, and de-crosslinked at 68°C for 14 hours. The DNA content was recovered using phenol-chloroform separation, and resuspended in 20 μl of 10 mM Tris-HCl, pH 8.0. For quantitative and allele-specific analysis, 7.6 μl of the final product was mixed with 8 μl of 2x QuantStudio 3D Digital PCR Master Mix v2 (Thermo A26358) and 0.4 μl Taqman SNP primer for rs6475604 (C 1754703_10) or rs1333040 (C 8766795_10) for negative control. The mixture was loaded onto the QuantStudio 3D Digital PCR system (Thermo) and analyzed using the provided cloud analysis system, following the manufacturer’s instructions.

### Luciferase reporter Assay

The enhancer region harboring rs6475604 (chr9: 22050310-22054084, hg38) was amplified from human mixed DNA (Promega G3041) using the following primer pairs: Fw: AAATCGATAAGGATCCAGGCCTTGCAATTGATTACG, Rev: AAGGGCATCGGTCGACGCACAAGAAATGCTAGCTAAGG, and cloned into the pGL4.23 backbone (Promega E8411) at the BamHI/SalI site using the In-Fusion cloning tool (Clontech). The product was transformed into Stellar competent cells (Clontech) to select clones with successful integration. After confirming the sequence using Sanger sequencing, desired clones were purified using the PureLink HiPure Plasmid Midiprep Kit (ThermoFisher). Five hundred nanograms of vector were co-transfected with 25 ng CMV-Renilla (Promega E2261) into subject cell lines in 24-well plates at 70% confluence using Lipofectamine 3000 reagent. After 24 hours, cells were analyzed using the Dual-Luciferase Reporter Assay System (Promega E1960) according to the manufacturer’s protocol. The reporter activity fold change was calculated with dual normalization against the empty pGL vector and Renilla reading. Luminescence was measured on a microplate luminometer (SpectraMax, Molecular Devices).

## Author Contribution

Conceptualization: YZ, YS; Methodology: YZ, CT, YS; Analysis: YZ, AF; Drafting: YZ, YS; Editing: all authors

---

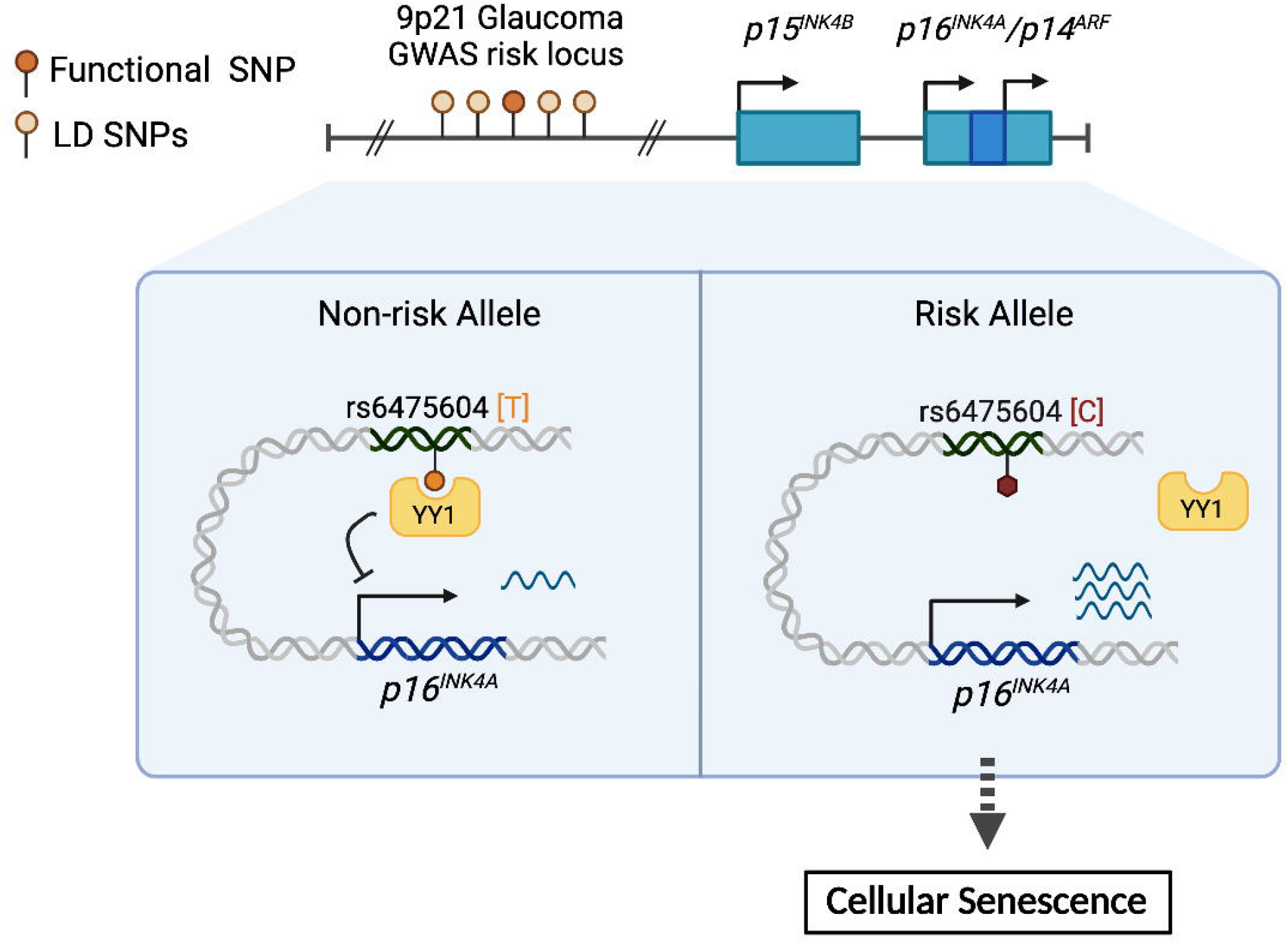

