## Supplementary material for "Identification and functional validation of an enhancer variant in the 9p21.3 locus associated with glaucoma risk and elevated expression of *p16^INK4a^*": Fig S1-S2; Table S1-S2

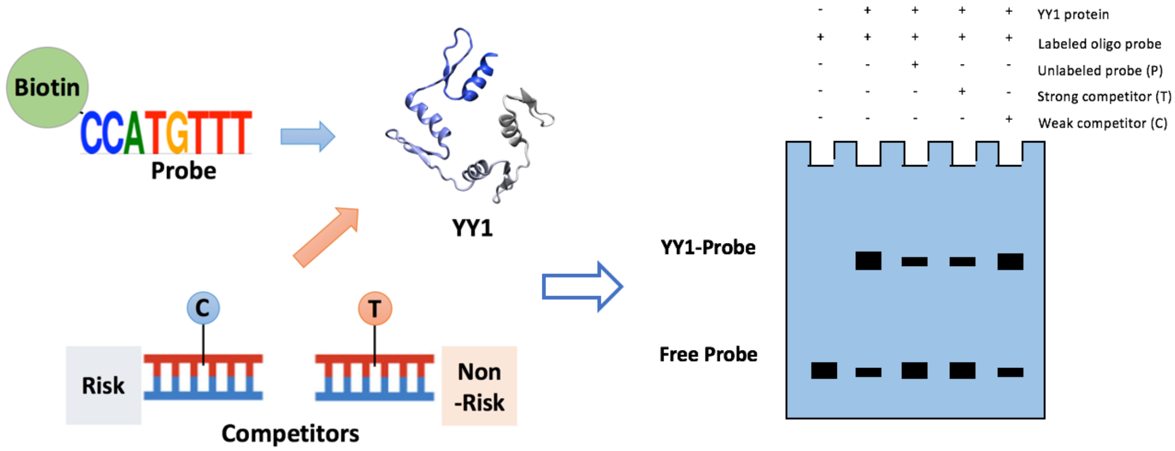


**Figure S1.** Schematic for the EMSA design and expected outcome from the competitive EMSA assay based on the computational prediction.

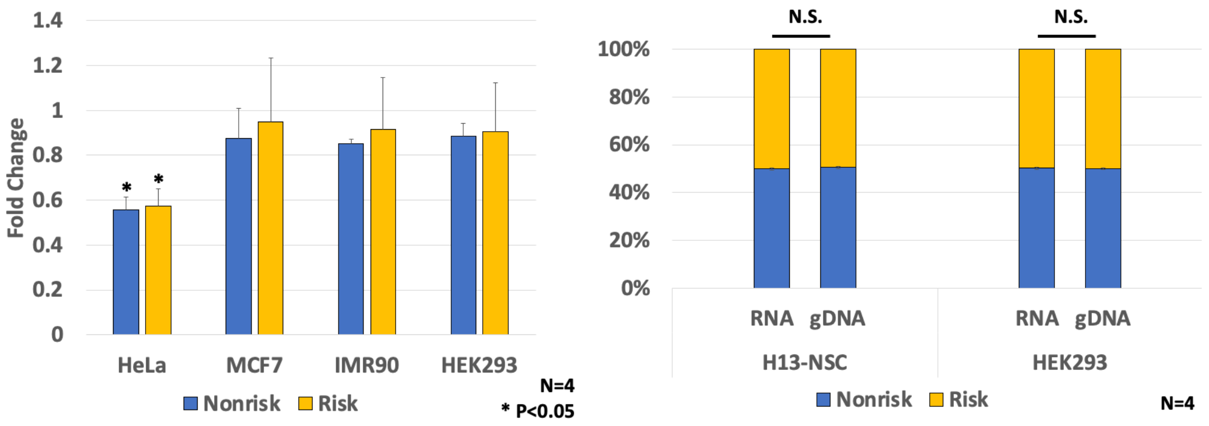


**Figure S2. (**A) Luciferase activity of rs6475604 enhancer region carrying non-risk or risk allele of the variant. (B) Allele-specific dPCR for CDKN2B expression in HEK293 and H13-derived neural stem cell.

| SNP ID | SNP POS | Ethinic Group | Location | Sample Size | p-value | Study |
|---|---|---|---|---|---|---|
| rs1063192-T | chr9:22003368 | Japanese | 3' UTR | 1394 Case, 6599 Controls | 5E-11 | Osman W |
| rs523096-A | chr9:22019130 | Japanese | Intronic | 286 Cases, 557 Controls | 5E-11 | Takamoto M |
| rs7865618-A | chr9:22031006 | Japanese | Intronic | 833 Cases, 686 Controls | 9E-11 | Nakano M |
| rs2157719-? | chr9:22033367 | European | Intronic | 3146 Cases, 3487 Controls | 2E-18 | Wiggs JL |
| rs1333037-T | chr9:22040766 | European | Intronic | 725 Cases, 11145 Controls | 1E-12 | Cooke Bailey JN |
| rs4977756-A | chr9:22068653 | European | Intronic | 1155 Cases, 1922 Contols | 7E-30 | Burdon KP Gharahkhani P |

**Table S1.** Reported GWAS variants associated with glaucoma in 9p21.3

| SNP ID | Chr | Pos | Reglome Score |
|---|---|---|---|
| rs6475604 | chr9 | 22052733 | 2a |
| rs3217977 | chr9 | 22007357 | 3a |
| **rs523096** | chr9 | 22019128 | 3a |
| **rs1333037** | chr9 | 22040764 | 4 |
| rs200059580 | chr9 | 22049479 | 4 |
| rs2069418 | chr9 | 22009697 | 4 |
| **rs1063192** | chr9 | 22003366 | 5 |
| rs10811648 | chr9 | 22067541 | 5 |
| rs10811649 | chr9 | 22067553 | 5 |
| rs10811651 | chr9 | 22067829 | 5 |
| rs10965224 | chr9 | 22067275 | 5 |
| rs1333039 | chr9 | 22065656 | 5 |
| rs1412829 | chr9 | 22043925 | 5 |
| rs142048183 | chr9 | 22028212 | 5 |
| **rs2157719** | chr9 | 22033365 | 5 |
| rs2383205 | chr9 | 22060934 | 5 |
| **rs4977756** | chr9 | 22068651 | 5 |
| rs581876 | chr9 | 22022375 | 5 |
| rs61271866 | chr9 | 21997014 | 5 |
| rs634537 | chr9 | 22032151 | 5 |
| rs679038 | chr9 | 22029079 | 5 |
| rs1008878 | chr9 | 22036111 | 6 |
| rs10965223 | chr9 | 22067003 | 6 |
| rs4451405 | chr9 | 22071749 | 6 |
| rs4977755 | chr9 | 22066362 | 6 |
| rs564398 | chr9 | 22029546 | 6 |
| rs599452 | chr9 | 22027401 | 6 |
| rs615552 | chr9 | 22026076 | 6 |
| rs7866783 | chr9 | 22056358 | 6 |
| rs1360589 | chr9 | 22045316 | 7 |
| rs1537378 | chr9 | 22061613 | 7 |
| rs1556515 | chr9 | 22036366 | 7 |
| rs2184061 | chr9 | 22061561 | 7 |
| rs518394 | chr9 | 22019672 | 7 |
| rs543830 | chr9 | 22026638 | 7 |
| rs613312 | chr9 | 22026593 | 7 |
| rs7030641 | chr9 | 22054039 | 7 |
| **rs7865618** | chr9 | 22031004 | 7 |
| rs8181050 | chr9 | 22064390 | 7 |
| rs944801 | chr9 | 22051669 | 7 |

**Table S2.** RegulomeDB score of 9p21 glaucoma risk variants ($r^2 > 0.8$). Score 7 is given to variants no available annotation data.
